# Adapting and optimizing GCaMP8f for use in *Caenorhabditis elegans*

**DOI:** 10.1101/2024.01.29.577882

**Authors:** Jun Liu, Elsa Bonnard, Monika Scholz

## Abstract

Improved genetically-encoded calcium indicators (GECIs) are essential for capturing intracellular dynamics of both muscle and neurons. A novel set of GECIs with ultra-fast kinetics and high sensitivity was recently reported by Zhang et al. (Nature, 2023). While these indicators, called jGCaMP8, were demonstrated to work in *Drosophila* and mice, data for *Caenorhabditis elegans* was not reported. Here, we present an optimized plasmid for *C. elegans* and use this to generate several strains expressing GCaMP8f. Utilizing the *myo-2* promoter, we compare pharyngeal muscle activity measured with GCaMP7f and GCaMP8f and find that GCaMP8f is brighter, shows faster kinetics and is less disruptive to the intrinsic contraction dynamics of the pharynx. Additionally, we validate its application for detecting neuronal activity in touch receptor neurons which reveals robust calcium transients at 25 ms time resolution. As such, we establish GCaMP8f as a potent tool for *C. elegans* research which is capable of extracting fast calcium dynamics at very low magnifications across multiple cell types.

## Introduction

Genetically encoded calcium indicators (GECIs) have been vital for studying the activity of many cell types including neurons (Inoue 2021), astrocytes (Lohr *et al*. 2021) and muscles (Kerr *et al*. 2000; Collins and Koelle 2013; Collins *et al*. 2016; Sanders *et al*. 2017), as well as their coordinated activity across the heart (Salgado-Almario *et al*. 2020) and whole brain (Ahrens and Engert 2015; Kato *et al*. 2015; Nguyen *et al*. 2016; Venkatachalam *et al*. 2016; Aimon *et al*. 2019; Hallinen *et al*. 2021; Markicevic *et al*. 2021; Atanas *et al*. 2022) using optical imaging. owing to this versatility, the past decades have seen rapid advancements in the brightness, photostability, sensitivity and binding kinetics of these indicators (Dana *et al*. 2019; Zhang *et al*. 2023), enabling experiments at faster timescales (Voleti *et al*. 2019) and with less disruption to normal animal physiology. A recent paper by Zhang et al., described a new generation of GECIs, derived from a prior generation of GECIs (GCaMP6) through structure-guided mutagenesis (Zhang *et al*. 2023). Named jGCaMP8, these indicators were demonstrated to have faster kinetics, to be brighter and more sensitive. While the study demonstrated these properties using recordings in the fruit fly, mouse and neuronal cell culture, its use in *C. elegans* was not demonstrated.

In *C. elegans*, GCaMP usage has been widespread and enabled discoveries about stimulus encoding of sensory neurons using single-cell imaging (Clark *et al*. 2006; Kato *et al*. 2014; Larsch *et al*. 2015; Itskovits *et al*. 2018; Katta *et al*. 2019) and multi-neuron imaging in chemosensory cells (Itskovits *et al*. 2018; Pritz *et al*. 2023). In addition, GCaMP was essential for detecting the compartment-specific activity in interneurons (Hendricks *et al*. 2012), and global effects of single neurons on activity and sleep (Turek *et al*. 2013). Going beyond single- or few neurons, whole-brain labeling even enabled the discovery of brain-wide coding of behavior in moving and restrained animals (Kato *et al*. 2015; Nguyen *et al*. 2016; Venkatachalam *et al*. 2016; Hallinen *et al*. 2021; Atanas *et al*. 2022).

With faster imaging technologies such as lightfield or lightsheet imaging (Voleti *et al*. 2019; Zhu *et al*. 2021) allowing brain imaging at up to 26 VPS, the true limitation for measurement is frequently the brightness of the indicator rather than the speed of acquisition (Kato *et al*. 2015; Hallinen *et al*. 2021; Atanas *et al*. 2022). Accordingly, faster and brighter indicators would positively impact these measurements in two ways, as they could either enable imaging of neurons driving faster behaviors, such as head swings, pharyngeal pumping and the activity of the spiking motoneurons (Gao and Zhen 2011; Hendricks *et al*. 2012; Trojanowski *et al*. 2016; Atanas *et al*. 2022), or alternatively enable longer measurements due to the lower photobleaching expected. We therefore adapted the published GCaMP8 for use in *C. elegans*, and measured its *in vivo* performance.

## Results

To determine if GCaMP8f also shows the reported improved properties as a fluorophore in *C. elegans*, we wanted to compare it with the prior generation of fluorophores in a standardized setting. GECIs are frequently compared in spiking neurons with stereotyped action potentials, which simplifies the analysis: as the underlying shape is expected to remain the same, any change in the signals read-out by imaging can be attributed to changes in the indicator properties. However, as *C. elegans* neurons predominantly used graded potentials (Goodman *et al*. 1998), we decided to instead use the stereotypical contractions of the pharynx to test the properties of this GECI. *C. elegans* uses its muscular pharynx to ingest bacteria, and this muscle shows action potentials shaped by voltage-gated calcium and potassium channels (Shtonda and Avery 2005). The quasi-rhythmic action of the pharyngeal muscle was also the first behavior whose activity was visualized with a GECI (i.e., Cameleon (Kerr *et al*. 2000)) in *C. elegans*.

To test the *C. elegans* optimized GCaMP8f, we used a similar approach and expressed the fluorophore in the pharyngeal muscle using the *myo-2* promoter. Based on the published sequence in (Zhang *et al*. 2023), we generated a codon-optimized version for *C. elegans*, with an additional three introns, that were suggested to improve expression (Okkema *et al*. 1993). The resulting sequence was cloned into a plasmid containing the *myo-2* promoter specific to pharyngeal muscle) and an *unc-54* 3’ UTR. We then compared the activity of GCaMP8f with the previous generation of GECI, GCaMP7f (Dana *et al*. 2019) (Fig. 1A,B). By applying serotonin (5-HT), action potentials in the pharynx can be stimulated and the activity of the GECI visualized using fluorescence microscopy with limited intrinsic behavioral variability (Fig. C). We find that as described for *Drosophila melanogaster*, mouse and in neuronal cell cultures (Zhang *et al*. 2023), the indicator is brighter (Fig. 1D, E) and shows faster kinetics compared to GCaMP7f (Fig. 1F).

**Fig. 1.**
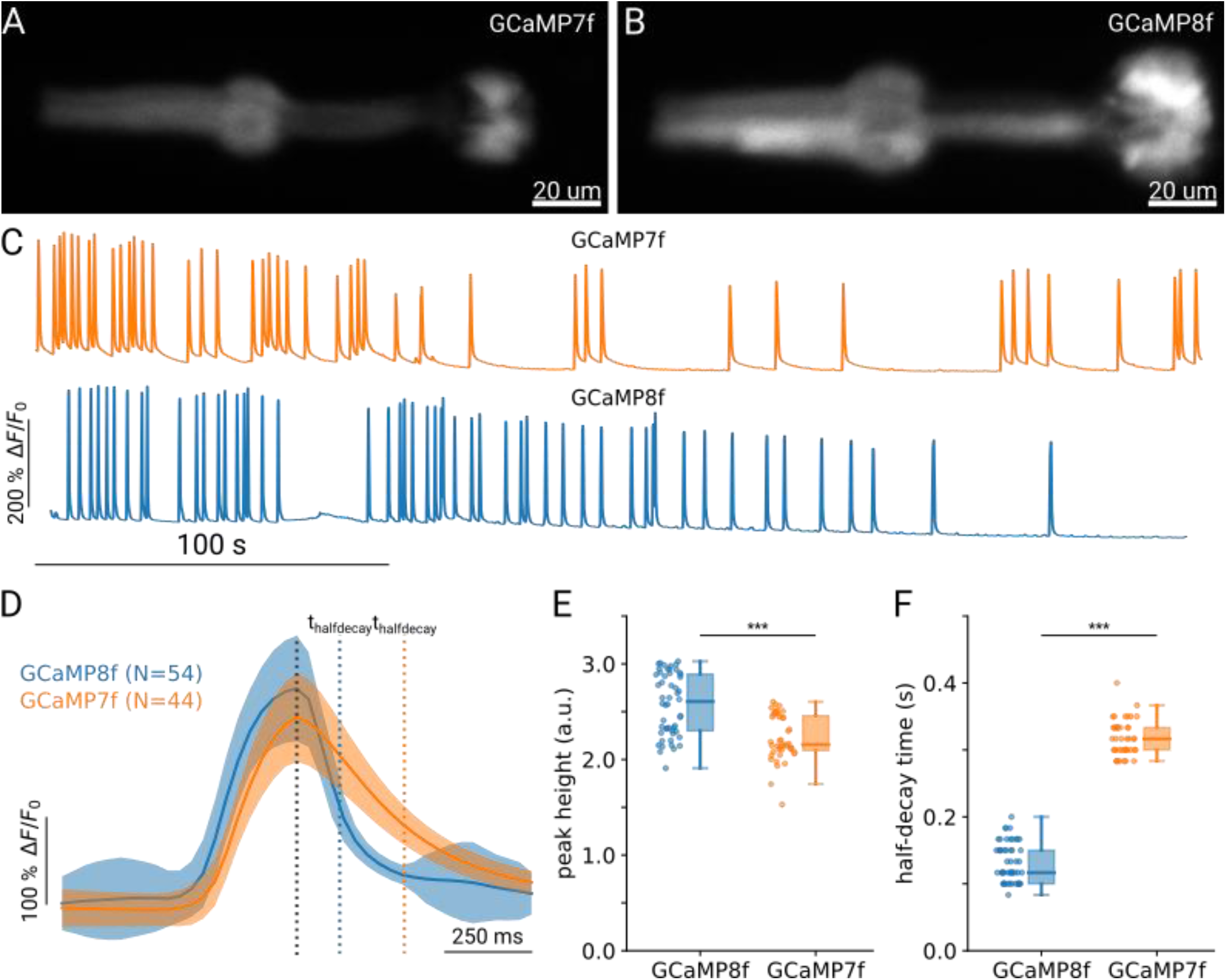
Improved detection of calcium dynamics with GCaMP8f due to faster indicator kinetics. (A) Images showing GCaMP7f and (B) GCaMP8f expressed under the *myo-2* promoter in adult *C. elegans* during pharyngeal pumping. (C) Example traces of pharyngeal muscle activity with GCaMP7f (orange) and GCaMP8f (blue) under stimulation with 10mM 5-HT. (D) Mean peak shape aligned to the maximal intensity for GCaMP7f (orange) and GCaMP8f (blue). The average half-decay time for each indicator is shown by the colored dashed lines. Shaded area indicates the standard deviation. (E) Maximal peak height for GCaMP7f (N=44) and GCaMP8f (N=54), respectively. (F) Half-decay time for each peak, as estimated from the first moment the traces reached 0.5*maximal height. *** indicates p-value <0.001. Significance was assessed using Welch’s unequal variance t-test.

Given the increased brightness, we wanted to test whether the speed and brightness enable new assays at lower resolution, allowing high-throughput data collection. While pumping can be detected without fluorescent labels, this requires imaging at higher magnifications and often laborious post-processing. We previously found that expressing a non-calcium dependent fluorophore enables detecting pumping at lower magnification and in freely moving animals (Bonnard *et al*. 2022). Using animals expressing either GCaMP7f or GCaMP8f in the pharyngeal muscle, we imaged groups of worms on a plate seeded with food. Compared to our previous work (Bonnard *et al*. 2022), we could detect animals labeled with GCaMP8f at even lower magnifications (0.5x instead of 1x), enabling a field of view of 1.5 cm (Fig. 2A, B). By tracking and segmenting the signals, we could find clear, stereotyped peaks in the fluorescence signals of animals expressing GCaMP8f, but not GCaMP7f (Fig. 2C). While the brightness of GCaMP7f was sufficient to track the animals, the peaks in GCaMP7f signals were noisy, likely due to the slower indicator speeds, and the lower brightness. We therefore concluded that the pumping rates extracted from peaks in GCaMP7f were not a reliable reflection of the true pumping rate (Fig.2E, gray boxplot). To verify the true pumping rate in the same conditions, we measured the pumping rate in brightfield images at high resolution for both strains, and found that the rate measured using GCaMP8f did not differ from the rate measured using high-resolution brightfield imaging Fig.2E. This indicator therefore enables robust, high-throughput analyses of feeding behavior which was not possible using GCaMP7f.

**Fig. 2.**
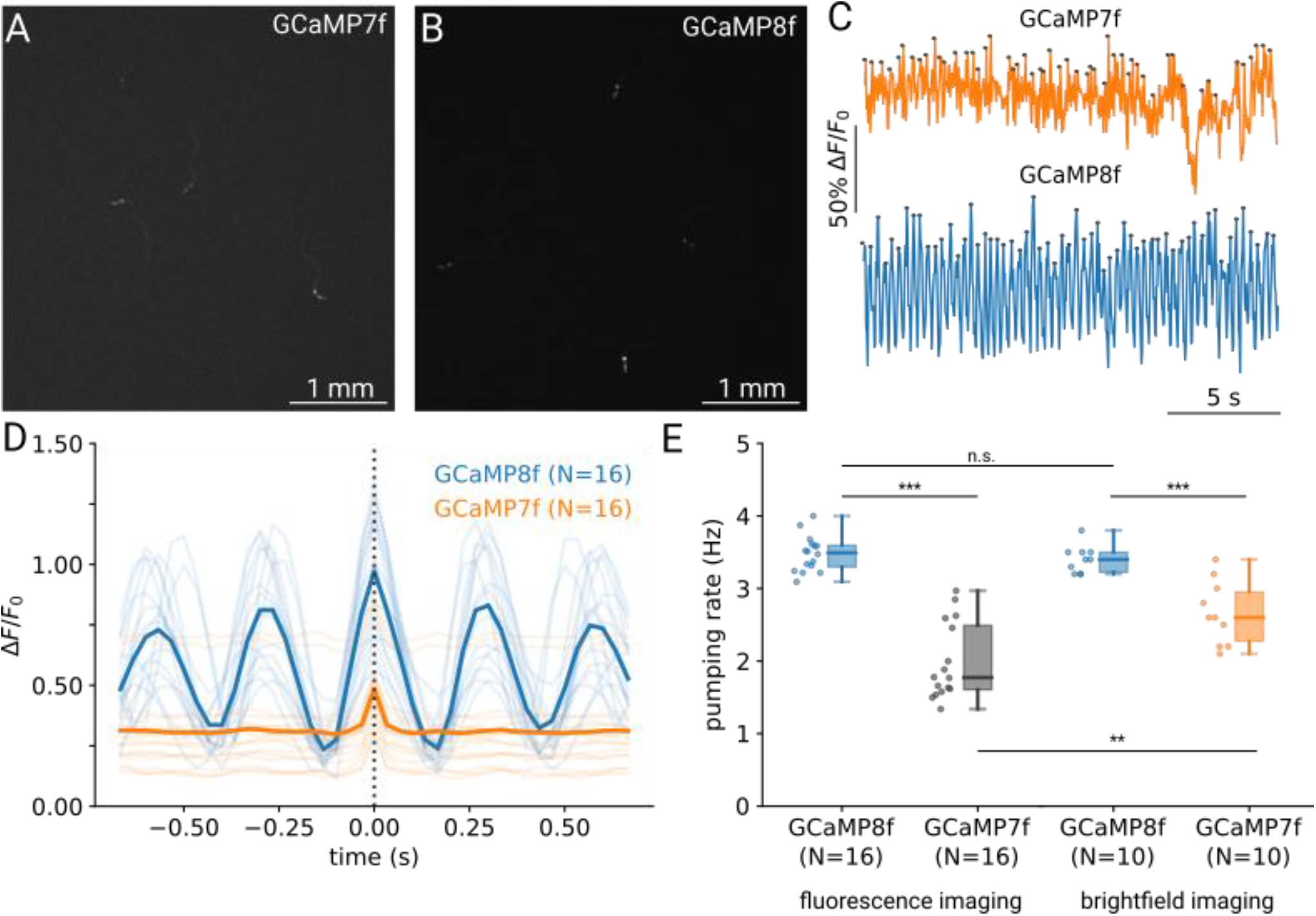
Large field-of-view measurements of pumping rates enabled by bright indicators. (A) Sample region showing animals expressing GCaMP7f and (B) GCaMP8f in the pharynx at low magnification of 0.5x. The full field-of-view comprises 14 mm x 10 mm. (C) Examples of resulting fluorescence activity traces for GCaMP7f (orange) and GCaMP8f (blue) with peaks indicated in gray. (D) Average of the detected peaks corresponding to pumping events for GCaMP7f (orange) and GCaMP8f (blue). (E) Pumping rate as measured from the peaks detected in (C) and by manual counting of brightfield images at higher resolution. Note that the results for GCaMP7f are only displayed for comparison, as the lower quality of the trace yields many false detections and does not result in reliable measurements. *** indicates p-value <0.001, ** indicates p-value<0.001. Significance was assessed using Welch’s unequal variance t-test and multiple comparisons were corrected using the Bonferroni-correction.

One disadvantage of these indicators is their required binding of intracellular calcium. This results in sequestering some of the free calcium ions, thereby lowering their overall concentration and potentially interfering with intracellular processes. The effects of this can be observed both for GECIs that express in muscle, as well as in neurons, as strains that express GCaMP pan-neuronally frequently move slower than wild-type animals (Nguyen *et al*. 2016; Yemini *et al*. 2021). Due to faster binding kinetics, we expect GCaMP8f animals to show less impairment than the animals expressing GCaMP7f, suggesting less interference with the calcium-dependent contractions. In line with these observations, we find that worms expressing GCaMP8f show significantly higher pumping rates on food compared to animals expressing GCaMP7f (Fig. 2D, E). We then compared the developmental speed of GCaMP7f and GCaMP8f worms by using the beginning of egg laying as a developmental reference. While all GCaMP8f worms had laid eggs within 73 h after being laid, only 5/8 GCaMP7f animals had laid their first eggs. Even after 77 hours, 3/8 animals had just reached their young adult stage, and not yet started laying eggs, supporting the observation that these animals developed less synchronously. This suggests the GCaMP8f is less disruptive to animal physiology than GCaMP7f.

We finally wanted to demonstrate that GCaMP8f also improves functional imaging of neuronal activity. As neurons are much smaller than the pharyngeal muscle, this typically requires higher magnifications, and yields noisier, lower amplitude signals compared to the data shown in Fig. 1. As touch receptor responses are also well characterized, they lend themselves to verify that GCaMP8f performs as expected and to check if this GECI allows imaging at faster frame rates or lower magnification. We therefore devised a protocol to stimulate the touch receptor neurons (targeting the TRNs using a *mec-17* promoter). We designed a plasmid containing both a baseline fluorophore (red, insensitive to intracellular calcium) and GCaMP8f. With this construct, multiple animals could be imaged simultaneously while a piezo buzzer supplied a gentle touch stimulus with varying amplitudes (Fig. 3A,B). By extracting the relative calcium changes, we could recapitulate the results of electrophysiological studies (O’Hagan *et al*. 2005; Eastwood *et al*. 2015; Katta *et al*. 2019) and imaging (Cho *et al*. 2017), demonstrating that TRN activity scales with stimulus amplitude (Fig. 3C,D). Compared to prior work using GCaMP6m the improved fluorophore GCaMP8f allowed us a faster imaging rate (40 Hz instead of 10 Hz) and at a lower magnification than usually used (20x instead of 40x or higher).

**Fig. 3.**
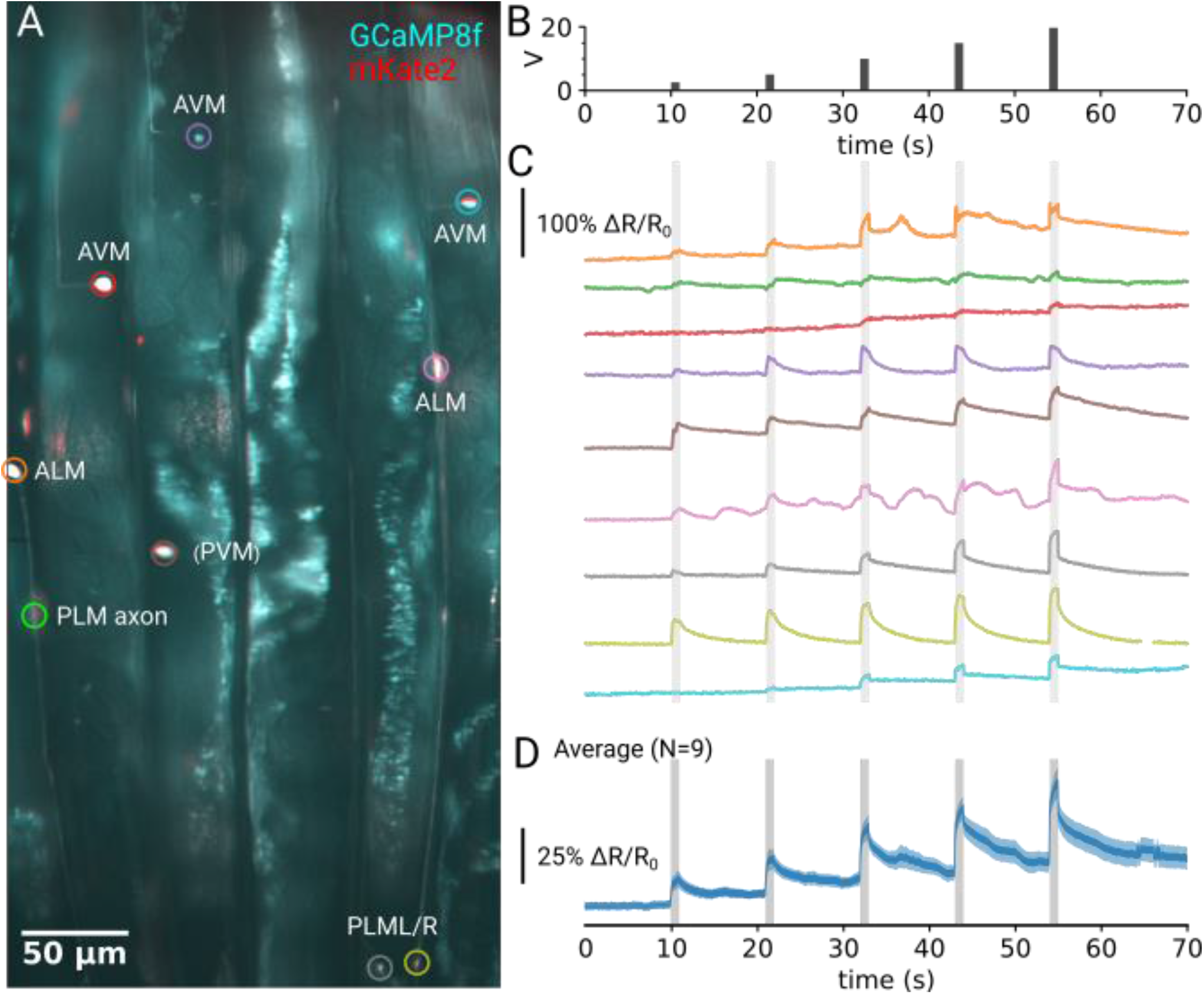
Rapid measurement of neural activity in response to gentle touch. (A) Overview of the field-of-view showing 5 animals arrayed longitudinally expressing GCaMP8f (cyan) and mKate2 (red, merged). The colored circles indicate regions used to extract Touch Receptor Neurons (TRNs) activity. (B) Stimulus profile applied to the sample using a 1s 630 Hz sinusoidal signal on a piezo buzzer. The traces correspond to the regions shown in (A). (C) Example neuronal traces using a ratiometric correction to reduce artifacts from motion or focus drift. (D) Average of the traces in (C) with the shaded area showing the s.e.m (N=9).

## Discussion

In summary, we find that in *C. elegans* GCaMP8f is brighter, faster and less disruptive to animal physiology than its immediate predecessor GCaMP7f. Accordingly, this improved fluorophore can enable new experiments studying subcellular compartments, activity of smaller, lower expression cells, or allow higher throughput experiments using more animals simultaneously by imaging at low magnifications (Larsch *et al*. 2013). Importantly, as GCaMP8f also appear to have a more limited impact on the nematode’s intrinsic biological processes, measurements using this fluorophore will be less disruptive and better reflect the dynamics and behavior of wild-type animals. Finally, for rapid whole-brain imaging, calcium indicators need to be either expressed at high levels or conversely, very bright to allow rapid imaging of a whole brain in 3D. Therefore, as imaging technologies have developed that allow faster 3D imaging, up to 26 VPS, in *C. elegans*, a new generation of fluorophore is required that keeps pace with these technological improvements. This generation of fluorophore will likely enable new experiments unlocking the neuronal coding of both behavior and stimuli on faster timescales.

## Methods

### Preparing imaging plates

For imaging pharyngeal muscle activity, imaging plates were prepared similar to standard nematode growth medium (NGM) plates, but without cholesterol, and agarose is used instead of agar to reduce autofluorescence.

For imaging the touch receptor neuron activity, an imaging chamber was prepared by filling a copper ring window (5 × 7 mm) by 2% agarose placed on an NGM plate. For ensuring the cohesion with the NGM block during the transfer to the stimulation apparatus, an extra 2% agarose on the copper ring edge was added.

### Imaging pharyngeal muscle activity at low resolution

30 adult animals were picked onto an imaging plate seeded with 150 ul of OP50 from an overnight culture. Animals were imaged on a commercial upright epi-fluorescence microscope (Axio Zoom V16; Zeiss) equipped with a 1 x objective (PlanNeoFluar Z 1.0 x/N.A. 0.25) with a camera sensor (acA3088-57um; BASLER) using a camera adapter with an additional 0.5 x magnification (60 N-C ⅔’’ 0.5 x; Zeiss) at 16 x nominal (0.5 x on camera) magnification resulting in a field of view of 1.5 cm x 1.0 cm. Videos of the lawn were recorded for 10 minutes at 30 fps. GCaMP7f images were recorded with a camera gain of 25, and GCaMP8f had to be reduced to a gain of 19 to avoid overexposure.

### Imaging pharyngeal muscle activity at high resolution

Adult animals were immersed in 20 ul of 10 mM 5-HT prepared with RediPrep™ Serotonin powder (InVivo Biosystems, USA) on a microscope slide. A coverslip was gently lowered onto the sample. The preparation was imaged on a commercial upright epi-fluorescence microscope (Axio Zoom V16; Zeiss) equipped with a 1 x objective (PlanNeoFluar Z 1.0 x/N.A. 0.25) with a camera sensor (acA3088-57um; BASLER) using a camera adapter with an additional 0.5 x magnification (60 N-C ⅔’’ 0.5 x; Zeiss) at 180 x nominal (5.626 x on camera) magnification. The resulting images were cropped to a region containing the entire pharynx of the worm using FIJI (Schindelin *et al*. 2012) and rotated for display. The relative fluorescence *ΔF/F*_0_ was calculated from the mean intensity across the entire image relative to the baseline (5 percentile of the data). Peaks were detected using python, and aligned to their respective maximum. The height of the peak and the half-decay time were extracted for each peak. Samples were selected for similar mean pumping rates, and lower mean rates.

### Analysis of pumping rates from brightfield images

Animals were imaged on a dissection microscope (Axio Zoom V16; Zeiss) at 125x nominal (0.614 um/px) resolution with a white LED light source. Videos were recorded for 30 s at 30 fps. Animals were manually kept centered in the field of view. 300 frames of data (corresponding to 10 s) that had a clearly visible grinder were counted by visualizing the data in Fiji (Schindelin *et al*. 2012). Pumping rates were calculated as pumping events/10s.

### Analysis of pharyngeal pumping rates in fluorescence images

Images were analyzed using Pharaglow (Bonnard *et al*. 2022). The peaks of the resulting intensity traces were found using the built-in peak detection and an average pumping rate per tracklet was calculated as N_pumps_/duration.

### Analysis of egg laying

Adult worms from INF418 (*nonEx106[myo-2p::GCaMP8f::unc-54 3’UTR]*) and INF96 (*syIs391[myo-2p::NLS::GAL4SK::VP64::unc-54 3’UTR + unc-122p::RFP + 1kb DNA ladder (NEB)]; syIs587[UAS::-GCaMP7f-SL2-mKate]*) were allowed to lay eggs for an hour to synchronize. Individual F1 worms were then transferred to separate plates to grow. For INF418, worms with and without extrachromosomal arrays were both picked, and the ones without the arrays were kept as non-transgenic siblings. The plates were checked once every 1-2 hours. The time of the first egg laying was recorded, except 3/8 plates from INF96, which appeared much younger and therefore the recording was discontinued when no eggs were detected after 77 h.

### Delivering touch stimulus

Substrate vibrations providing a gentle touch stimulus were delivered by gluing a 48 mm diameter piezo buzzer (APS4812B-LW100-R, PUI Audio Inc., USA) on a 60 mm diameter petri-dish housing the imaging chamber. The piezo element was driven by a 1-second 630 Hz sinusoidal voltage at various amplitudes (2, 5, 10, 15 and 20 Volts peak to peak (Vpp)). The stimulus delivery and the synchronization with the camera acquisition were controlled in a customized program (LabVIEW, National Instruments, USA) using a data acquisition card (BNC-2090A, National Instrument).

### Imaging touch receptor neuron activity

Six adult animals were picked onto the imaging chamber and immobilized in 10 mM Levamisol (Sigma-Aldrich). The animals were imaged using an epifluorescence microscope for ratiometric calcium imaging equipped with a 20 x objective (S Plan Fluor LWD, N.A. 0.70, Nikon). The resulting field of view was 250 × 500 um with a pixel size of 0.24 um per pixel.

To excite GCaMP8f and mKate2 fluorophores, the cyan (470/24 nm, Chroma) and green (575/15 nm, Semrock) lines from an LED lamp (Spectra X light engine, Lumencor) were projected onto the sample. Transmitted and emitted light were filtered using a triple-edge dichroic beamsplitter (409/493/596 nm, Semrock). To simultaneously image GCaMP8f and mKate2, a dual view with a 585 nm beamsplitter (DV2, Photometrics) was used. Each channel was projected onto a half of an sCMOS camera (Zyla, Andor) at 40 Hz acquisition rate with a 16 bit readout depth.

### Ratiometric analysis

The resulting images were automatically split into individual channel images and registered for an optimal overlay using Matlab. Using the mKate2 channel images, manually defined regions around each neuron including background were cropped in Fiji. Within each region, the neuron was tracked using the Fiji plugin TrackMate (Tinevez *et al*. 2017) defining a circular region of interest around the neuron. In Python, the fluorescent signal in the circular ROI in both channels images was extracted and the ratio change was calculated as follow:

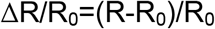

Where R is the ratio between GCaMP8f and mKate2 signals, and R_0_ the baseline ratio calculated from the time-averaged signals 10 s before the stimulus.

### Reagents

Plasmids: pLJ52 was adapted from the sequence described in (Zhang *et al*. 2023) by codon optimizing the GCaMP8f and adding 3 synthetic introns as described in (Redemann *et al*. 2011). The sequence was created using the tool at https://worm.mpi-cbg.de/codons/cgi-bin/optimize.py and synthesized by GenScript. Then GCaMP8f was subcloned into an expression vector (derived from pRL231, gift of Manuel Zimmer) to make pLJ53 (*myo-2p::GCaMP8f::unc-54 3’utr*).

“SL2::mKate2::let-858_3’UTR” was used to replace the *unc-54 3’UTR* in pLJ53 to make pLJ54 (*myo-2p_GCaMP8f_SL2_mKate2_let-858_3’UTR*). Then *myo-2p* was further replaced by *mec-17p* to make pLJ57 (*mec-17p_GCaMP8f_SL2_mKate2_let-858_3’UTR*).

| Strain Name | Genotype | Short Name in Article | Source |
| --- | --- | --- | --- |
| INF96 | *syls391[myo-2p::NLS::GAL4SK::VP64::unc-54 3'UTR + unc-122p::RFP + 1kb DNA ladder (NEB)]; syls587[UAS::GCaMP7f-SL2-mKate]* | GCaMP7f | Cross of PS7154 to PS8423 from (Wang *et al.* 2017) |
| INF418 | *nonEx106[myo-2p::GCaMP8f::unc-54 3'UTR]* | GCaMP8f | This study. pLJ53 was injected into N2 at 5ng/ul. |
| INF447 | *nonEx133[mec-17p_GCaMP8f_SL2_mKate2_let-858_3'UTR + Punc-122::tagBFP]* | TRN::GCaMP8f | This study. pLJ57 was injected into N2 at 40ng/ul. *coel::bfp* (pLJ49) was used as a marker (40ng/ul). |

## Data availability statement

Strains and plasmids are available upon request for all strains and plasmids generated in this study. The plasmid maps for pLJ53 and pLJ57 are available as File S1 and File S2, respectively. All data underlying the figures will be made available on OSF after publication. A reviewer-only link is provided here: https://osf.io/4pw2m/?view_only=53f8dfc2cd174157819054a2ffe1d1b4

## Acknowledgements

We thank Paul Sternberg for sharing the cGAL strains, especially the myo-2p driver and GCaMP7f effector lines with us. We also thank Manuel Zimmer for sharing the pRL231 plasmid. We thank James Lightfoot for helpful discussions and comments on this manuscript. Some strains were provided by the CGC, which is funded by NIH Office of Research Infrastructure Programs (P40 OD010440). This work was funded by the Max Planck Society.

